# Making sense of touch: spatial tactile cues promote efficient visuo–somatosensory integration and restore motor accuracy under sensory conflict

**DOI:** 10.64898/2026.02.10.704673

**Authors:** Maria Evangelia Vlachou, Eva Lafaverges, Laurence Mouchnino, Jean Blouin

## Abstract

Effective interaction with complex environments relies on integrating somatosensory and visual signals into coherent spatial representations of the body in external space. When these signals conflict, as during movement under mirror-reversed visual feedback, motor performance deteriorates. Under such conflict, visual information typically facilitates motor performance, whereas tactile feedback impairs it. Here, we tested whether providing tactile input with explicit spatial structure renders visuo-somatosensory integration functionally relevant and stabilizes motor behaviour under sensory conflict. Healthy adults performed a mirror-tracing task, tracing a shape’s contour with the index finger while tactile feedback conveyed either motion-related cues alone or additional discrete spatial landmarks aligned with the outline. Spatially structured tactile cues preserved accuracy and smoothness relative to motion-related tactile input, indicating more effective online motor control. EEG source analyses further revealed that, when spatial tactile cues were present, somatosensory activity increased and functional connectivity became more focal within sensorimotor networks. In contrast, when absent, sensory conflict was associated with widespread reductions in cortical activity, including in somatosensory areas, and with a diffuse functional connectivity pattern consistent with elevated sensorimotor uncertainty. Together, these findings demonstrate that spatial tactile cues bias hand-displacement encoding toward an external reference frame, promote efficient visuo– somatosensory integration and mitigate the detrimental impact of sensory conflict on motor action.

## Introduction

Tactile signals arising from mechanoreceptors ascend via the dorsal column-medial lemniscus pathway to the primary somatosensory cortex (S1), where they undergo initial cortical processing. Within S1, tactile inputs are organized somatotopically, providing a body-surface-centred reference frame that supports precise localization of skin stimulation (for a review, Delhaye et al., 2018)^1^. Through integration with proprioceptive and visual signals, together with predictions derived from efference copies, this local cutaneous map contributes to the formation of egocentric representations of limb position^2–4^. During interaction with external objects and surfaces, however, tactile information can be transformed into allocentric, environment-based coordinates^5–7^, extending its functional relevance to perception and action. Converging evidence implicates the posterior parietal cortex (PPC) as a key site for these multisensory transformations and spatial remapping processes^8–10^.

While substantial work has examined how tactile information is perceptually encoded across reference frames, far less is known about how egocentric and allocentric coding of spatial touch supports movement control. Isolating the specific contribution of tactile feedback during visually guided actions is challenging because visual information often induces rapid ceiling effects: high movement accuracy can be achieved through vision alone, thereby masking potential influences of tactile signals. To address this limitation, we recently examined hand kinematics and somatosensory cortical activity while participants traced the outline of a polygon with the pulp of their index finger under mirror-reversed visual feedback^11^. Under such condition, analogous to those encountered in video-assisted surgery or microscope-guided manipulation, visual and somatosensory feedback become spatially incongruent. A hallmark of this sensory context is that processing somatosensory feedback from the hand impairs movement control^12,13^. By comparing performance under intact tactile input with a condition in which tactile feedback was reduced using a finger splint, we tested whether tactile feedback contributes to motor control despite visual dominance. Crucially, we found that tracing performance deteriorated more when tactile feedback was intact than when it was attenuated by the splint^11^, consistent with the hypothesis that tactile cues are indeed integrated within the sensorimotor loop, even in the presence of visual feedback. However, because somatosensory input is irrelevant for controlling movement under mirror vision, its processing disrupts performance. Under full cutaneous feedback, and in line with sensory weighting theory, somatosensory cortical activity was reduced, suggesting a down-weighting of tactile information that conflicted with visual input and impaired movement control under mirror vision. In contrast, no such gating occurred when tactile signals were already reduced by the finger splint.

Building on evidence that tactile edges convey spatial information supporting accurate localization and orientation in allocentric (object- or environment-centred) reference frames^14^, and on findings that allocentric encoding of environmental and movement-related cues improves performance under sensory conflict^13,15^, we hypothesized that providing explicit spatial tactile information would facilitate motor performance when visual and somatosensory feedback are misaligned. Accordingly, we predicted that the reduction in somatosensory processing typically observed under visuo-somatosensory conflict^12,13,16,17^ should be attenuated, or even reversed, when spatially structured tactile cues are available during tracing.

To test this prediction, we adapted the mirror-tracing paradigm to manipulate the spatial content of tactile feedback (Fig. 1a). One group (Tactile group) traced the shape on a heterogeneously textured surface in which the outline, composed of Braille-like points, differed from the smooth background in both colour and texture (Fig. 1b). In this configuration, tactile input conveyed not only motion-related information arising from finger-surface interactions but also discrete spatial landmarks that could anchor touch to stable external structures. A second group (Visual group) performed the same task on a homogeneously textured surface, where the outline differed from the background only in colour (Fig. 1c). In this case, tactile input was limited to motion-related cues and lacked explicit features that could serve as spatial landmarks for spatiotopic interpretation.

**Fig. 1.**
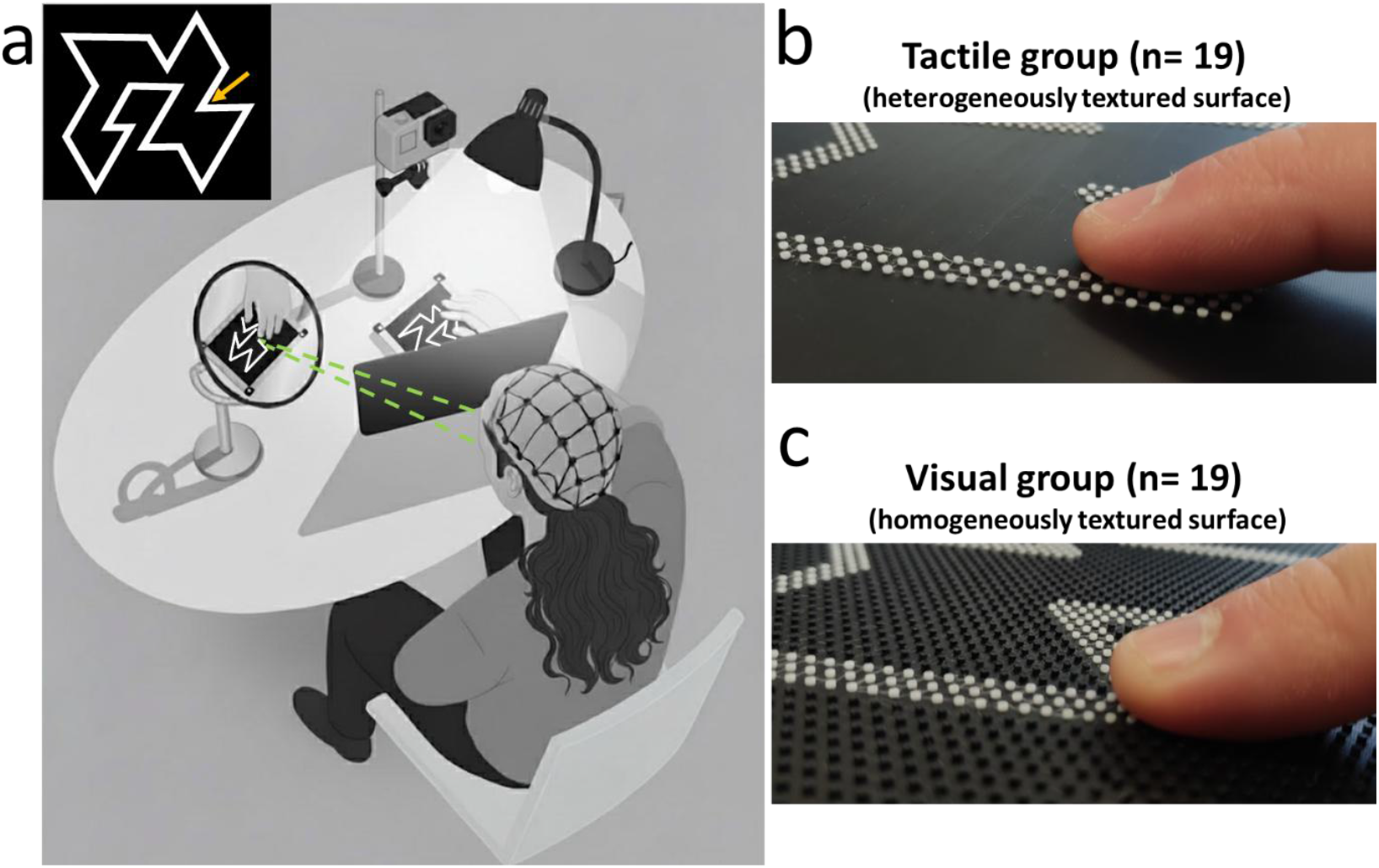
Mirror-tracing task design. **a** Healthy young adults traced the outline of the irregular polygon on a surface screwed on an AMTI force platform, while looking at the reflection of their hand and the polygon in an angled mirror. A shield prevented direct vision of both the hand and the tracing surface. Kinematics, contact forces and EEG activity were recorded throughout the task. In a control condition, participants performed the same tracing task with direct visual guidance (not illustrated). The inset shows the polygon used in the experiment; the yellow arrow indicates the starting point for each session (for illustration purposes only and not visible on the tracing surface). After each trial, participants resumed tracing from the point at which they had stopped on the previous trial. **b** Close-up of the finger–surface interaction in the Tactile group. Participants traced the polygon on a heterogeneously textured surface in which the outline was distinguished from the background by both colour (white vs. black) and texture (raised dotted pattern vs. smooth background). **c** Close-up of the finger–surface interaction in the Visual group. Participants traced the same polygon on a homogeneously textured dotted surface, in which the white outline differed from the black background only in colour.

## Results

Detailed statistical outputs for all reported analyses are available in the Supplementary material.

### Explicit spatial tactile cues preserved tracing accuracy during sensory conflict

We evaluated task performance by examining two key aspects of tracing: accuracy and smoothness. Tracing accuracy was quantified using the distance-to-segment index (hereafter referred to as the error index), a well-established metric for assessing movement precision under sensory conflict^11–13,18–21^. This index, derived from video tracking data, represents the ratio between the total distance traced by the participant’s finger and the actual length of the corresponding polygon segment. A value of 1 indicates optimal performance (the traced distance matched the actual path length), while higher ratios reflect greater deviations from the intended path and thus reduced tracing accuracy (greater error).

Figure 2a illustrates the median and distribution of the error index for both the Tactile and Visual groups during the Direct (control) and Mirror vision conditions. A 2 (Group: Tactile, Visual) × 2 (Condition: Direct vision, Mirror vision) mixed ANOVA revealed a significant Group × Condition interaction (F_1,35_=9.2, p = .005, η^2^_p_ = 0.21). Post hoc pairwise comparisons using Fisher’s LSD test indicated that the error index increased significantly under mirror-reversed relative to direct vision in the Visual group (p < .001, Cohen’s d = −2.158), but did not significantly change across visual conditions in the Tactile group (p = .19).

**Fig. 2.**
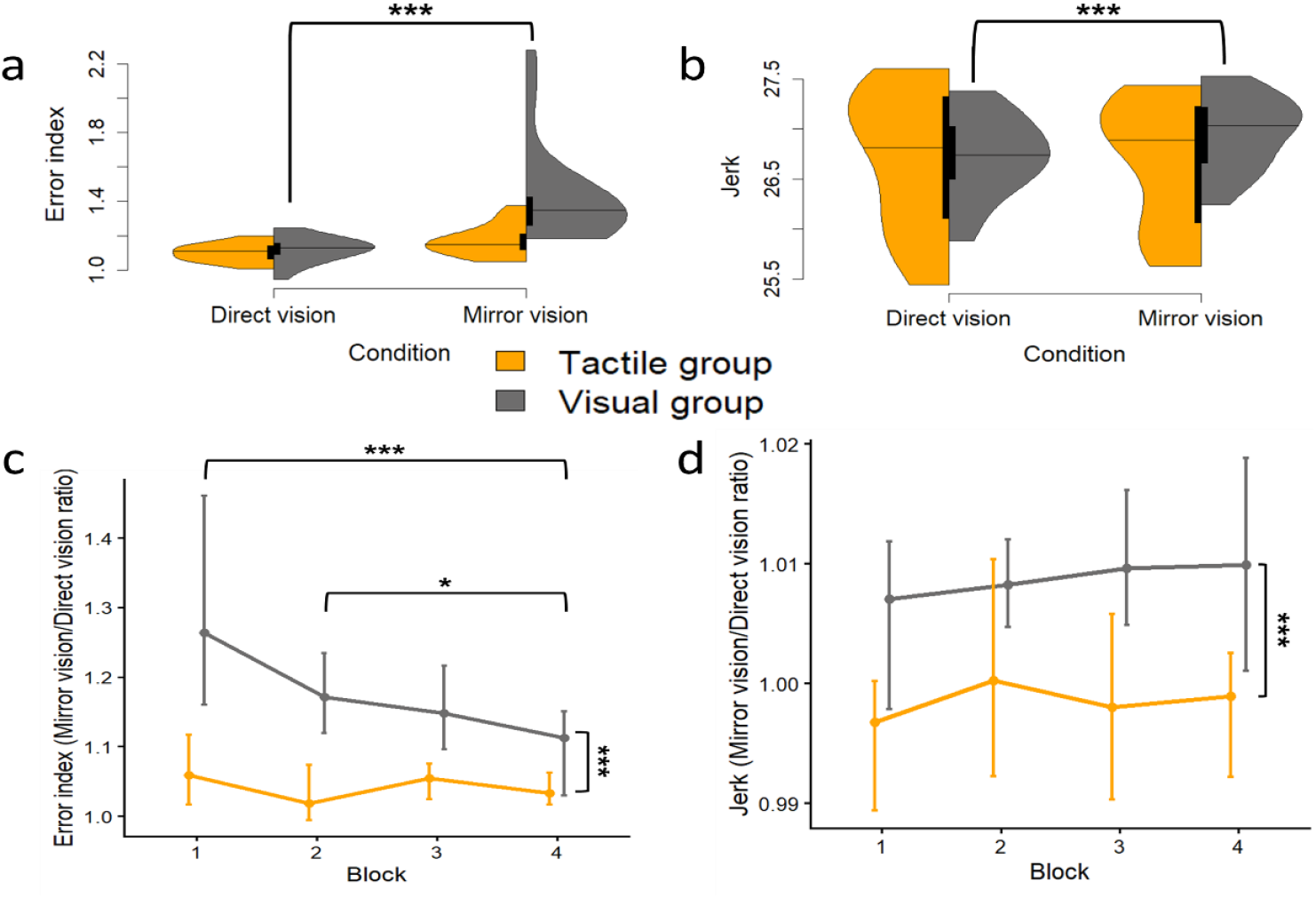
Spatial tactile cues preserve movement accuracy and smoothness during mirror tracing. **a, b** Half violin plots showing the distribution of the median error **(a)** and jerk **(b)** indices across all trials, for both the Tactile (orange) and Visual (grey) groups under Mirror and Direct vision conditions. Black vertical boxes represent the interquartile range (IQR; middle 50% of the data) and horizontal lines indicate the median. **c, d** Line plots depicting the evolution, across 5-trial blocks and for both the Tactile (orange) and Visual (grey) groups, of the **(c)** median error index ratio (Mirror/Direct vision condition) and **(d)** jerk index ratio (Mirror/Direct vision condition). A ratio of 1 represents optimal tracing performance. Error bars represent the ±95% confidence interval of the median. * p < .05, *** p < .001.

Movement smoothness captures the temporal continuity and efficiency of motor execution, which has been related to effort minimization and optimal spatiotemporal coordination^22,23^. It was quantified by using the log dimensionless jerk (hereafter referred to as jerk), computed from the rate of change of the resultant fingertip tangential force in the XY plane of the surface. The jerk therefore reflects the abruptness of force transitions, providing an empirical measure of movement coordination and stability. A 2 (Group: Tactile, Visual) × 2 (Condition: Direct vision, Mirror vision) mixed ANOVA on jerk revealed a significant Group x Condition interaction effect (F_1,35_=13.89, p < .001, η^2^_p_ = 0.28) (Fig. 2b). Similarly to the pattern observed for the accuracy index, mirror-reversed vision led to higher jerk values in the Visual group (p < .001, Cohen’s d = −0.426), while jerk remained unaffected by visual condition in the Tactile group (p = .33).

Several procedural measures were implemented to minimize sensory adaptation (see Methods) and to increase the likelihood that EEG activity reflected stable sensorimotor processing across the session, thereby allowing the averaging of EEG data across trials. To evaluate potential adaptation effects behaviourally, we examined changes in the error and jerk indices over the course of the experimental session. The 20 trials were grouped into four consecutive blocks of five trials, and for each block, we computed the mean ratio between Mirror and Direct vision performance (Fig. 2c, d). Because the distribution of the error-ratio data violated parametric assumptions, non-parametric Friedman tests were conducted separately for each group. In the Tactile group, no significant effect of trial block was observed (p = .926), indicating stable performance across the session. In contrast, the Visual group showed a significant effect of trial block (p = .002). Conover’s post hoc analysis revealed that the error index ratio in the first (p < .001, r_rb_ = 0.8) and second (p = .037, r_rb_ = 0.71) blocks was significantly higher than in the last block, reflecting improved performance over time. However, the Visual group exhibited significantly higher error indices than the Tactile group in all trial blocks, as confirmed by non-parametric Mann–Whitney tests conducted for each block (1–5: p = .04, r_rb_ = 0.54; 6–10: p < .001, r_rb_ = 0.69; 11–15: p < .001, r_rb_ = 0.62; 16–20: p = .031, r_rb_ = 0.42).

The evolution of jerk ratio (Mirror vision/Direct vision condition) over time is shown in Figure 2d. A 2 (Group: Tactile, Visual) × 4 (Block: 1, 2, 3, 4) mixed ANOVA revealed a significant Group effect (F_1,35_=13.4, p < .001, η^2^_p_ = 0.28) with the Visual group exhibiting higher values. No significant effect of Block or Group × Block interaction was observed, indicating a stable movement smoothness profile for both groups across the session.

Taken together, the availability of explicit spatial tactile cues preserved tracing performance under sensory conflict, yielding greater movement accuracy and smoothness than when tactile input lacked external spatial structure. Although tracing accuracy in the Visual group improved over time, it never reached the level achieved by the Tactile group. This persistent performance gap supports the rationale for analysing and comparing EEG activity averaged across all trials in the following section.

### Increased somatosensory cortical activity with spatial tactile cues

All EEG analyses were conducted in source space (see Methods), allowing us to examine how spatial tactile cues modulated brain activity, measured as absolute current amplitude, within anatomically and functionally defined regions during tracing performed under visuo-somatosensory conflict^24^. Specifically, we assessed within-group effects of sensory conflict by contrasting EEG current in the Mirror vision condition with the Direct vision (control) condition for each experimental group. Statistical comparisons were performed using two-tailed paired permutation Wilcoxon signed-rank tests (significance threshold: p < 0.05, uncorrected) applied to the mean current over the 24-second tracing period.

The statistical topographical current maps in Fig. 3 depict the spatial distribution of condition-dependent cortical activation. Warm colours indicate regions where mirror-reversed visual feedback elicited greater cortical activity relative to direct vision, while cool colours represent regions of reduced activation under mirror vision. Grey areas correspond to regions showing no significant differences between conditions. Within sensory cortices, warm colours are interpreted as reflecting sensory facilitation, consistent with enhanced afferent processing or increased engagement in response to altered visual feedback. Conversely, cool colours are indicative of sensory gating, consistent with reduced transmission of afferent feedback.

**Fig. 3.**
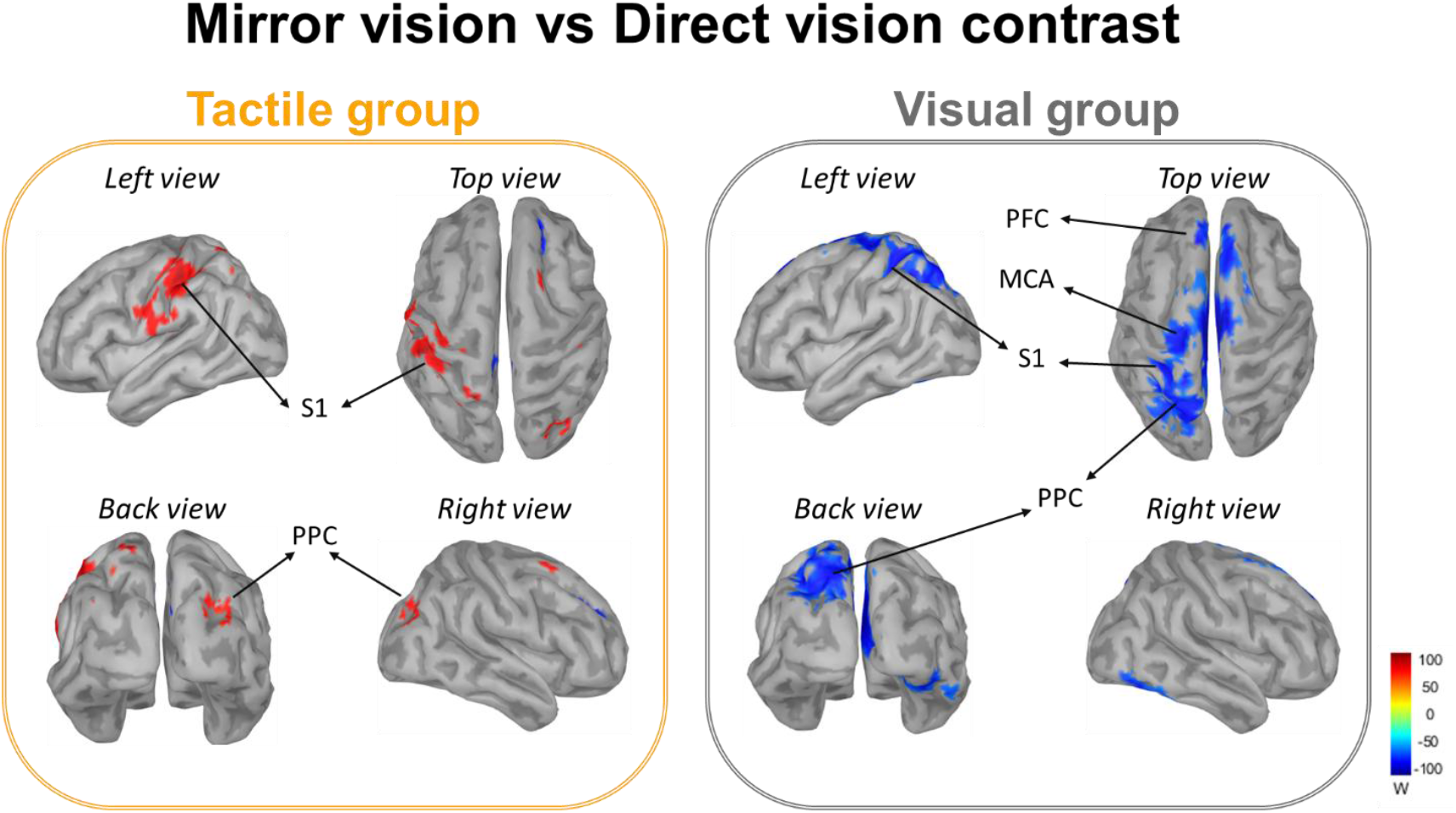
Spatial information in tactile cues reshape cortical activation patterns. Statistical topographical current maps resulting from the contrast between the Mirror and Direct vision conditions for the Tactile (left) and the Visual groups (right). Source estimates were projected onto a standard cortical template (MNI Colin27; 15,002 vertices). For each group, left and right lateral views, as well as back and top views, are displayed. MCA: motor cortex areas; PFC: prefrontal cortex; PPC: posterior parietal cortex; S1: primary somatosensory cortex.

The impact of visuo-somatosensory conflict on cortical activity was strongly modulated by the spatial content of tactile cues during tracing, as evidenced by significant within-group effects. When the surface dots conveyed information about the shape’s outline (Tactile group), Mirror vision elicited increased activity in the left primary somatosensory (S1) and right posterior parietal (PPC) cortices relative to the Direct vision (control) condition (Fig. 3, left panel). By contrast, when tracing was performed on the homogeneously, no-spatially informative textured surface (Visual group), mirror vision led to a widespread reduction in cortical activity relative to the Direct vision condition, encompassing S1, PPC, motor areas and prefrontal cortex (Fig. 3, right panel).

Given our focus on how spatial tactile information modulates somatosensory processing, we next examined group differences in activity within key somatosensory and parietal regions. Mean absolute source amplitudes were extracted from predefined regions of interest (ROIs) in the left hemisphere, contralateral to the tracing hand, encompassing the postcentral gyrus (S1), anterior superior parietal lobule (aSPL) and posterior superior parietal lobule (pSPL). These ROI-specific source amplitudes were then submitted to separate 2 (Group: Tactile, Visual) × 2 (Condition: Direct vision, Mirror vision) mixed-design ANOVAs, with Condition as the within-subject factor (Fig. 4).

**Fig. 4.**
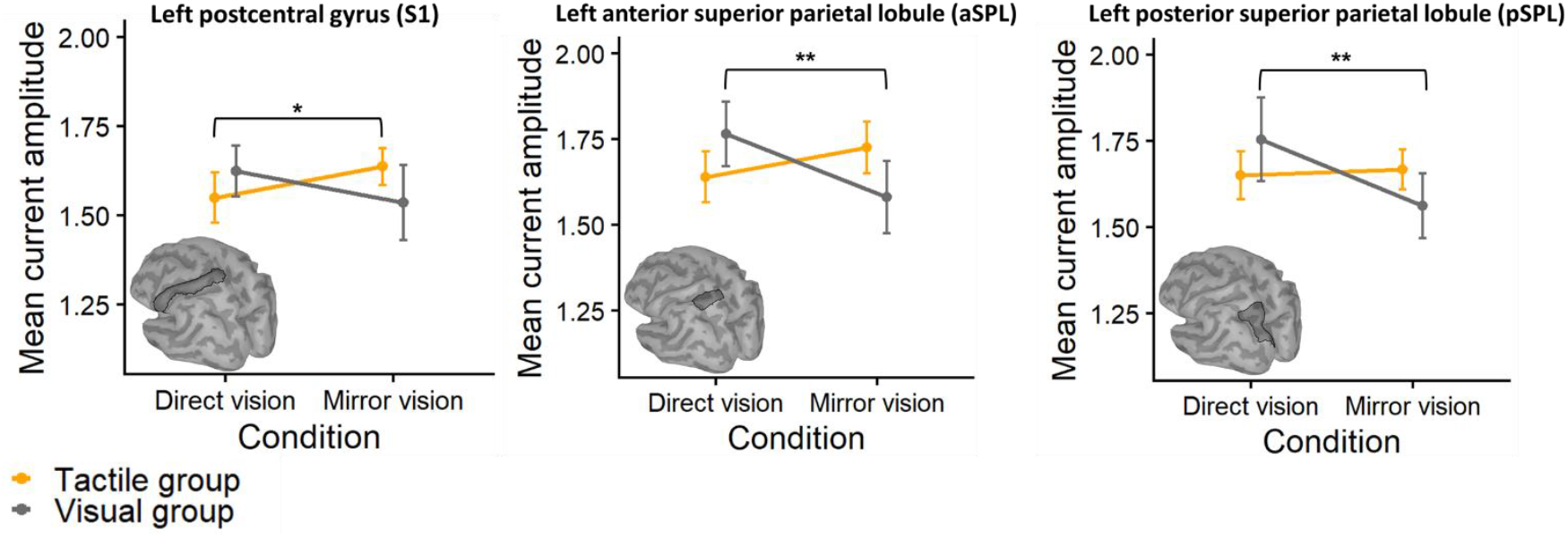
Effect of tactile cue relevance on somatosensory cortical activity. Line plots show the mean absolute source current amplitude in predefined regions of interest (ROIs) for the Tactile and Visual groups under Direct vision and Mirror vision conditions. ROIs include the left postcentral gyrus primary somatosensory cortex (S1, left), anterior superior parietal lobule (aSPL, middle), and posterior superior parietal lobule (pSPL, right). Data reveal region- and group-specific modulations of cortical activity in response to mirror-reversed visual feedback. Error bars represent the ±95% confidence interval (CI) of the mean. *p < .05, **p < .01.

For the S1 ROI, the ANOVA revealed a significant Group × Condition interaction (F_2_,_29_ = 6.9, p = .01, η^2^_p_ = 0.19). Fisher’s LSD post hoc comparisons further revealed that current amplitude increased significantly in the Tactile group under Mirror vision relative to Direct vision condition (p =.01, Cohen’s d = -0.9), whereas no significant difference was observed in the Visual group (p=.32). Similarly, significant Group × Condition interactions were observed for both the aSPL ROI (F_2, 30_ = 10.42, p = .003, η^2^_p_ = 0.26) and the pSPL ROI (F_2_,_30_ = 7.45, p = .011, η^2^_p_ = 0.2). Post hoc analyses indicated that, in the Visual group, current amplitudes decreased significantly under Mirror vision relative to the Direct vision condition in both regions (aSPL: p = .004, Cohen’s d = 0.43; pSPL: p = .0013, Cohen’s d = 0.23), whereas no significant changes were observed in the Tactile group (aSPL: p = .16; pSPL: p = .76).

These findings indicate that mirror-reversed visual feedback modulated somatosensory cortical activity in a region- and group-specific manner: the Tactile group showed enhanced S1 activity, whereas Visual group exhibited suppression in posterior parietal areas involved in somatosensory integration.

Finally, we examined how task-relevant spatial tactile inputs influenced functional connectivity among cortical ROIs implicated in visually guided tracing, visuo-somatosensory integration, and sensory conflict resolution^25–28^. These bilateral, non-overlapping ROIs included the S1, aSPL, pSPL, lateral occipital cortex (LOC), supramarginal gyrus (SMG), prefrontal cortex (PFC), premotor cortex (PMC), and the supplementary motor area (SMA) (Fig. 5). For each group, we computed a non-directed connectivity metric based on temporal associations between the ROI time series and compared the Mirror and Direct vision conditions using a two-tailed paired permutation Wilcoxon signed-rank test (significance threshold p < .05, uncorrected).

**Fig. 5.**
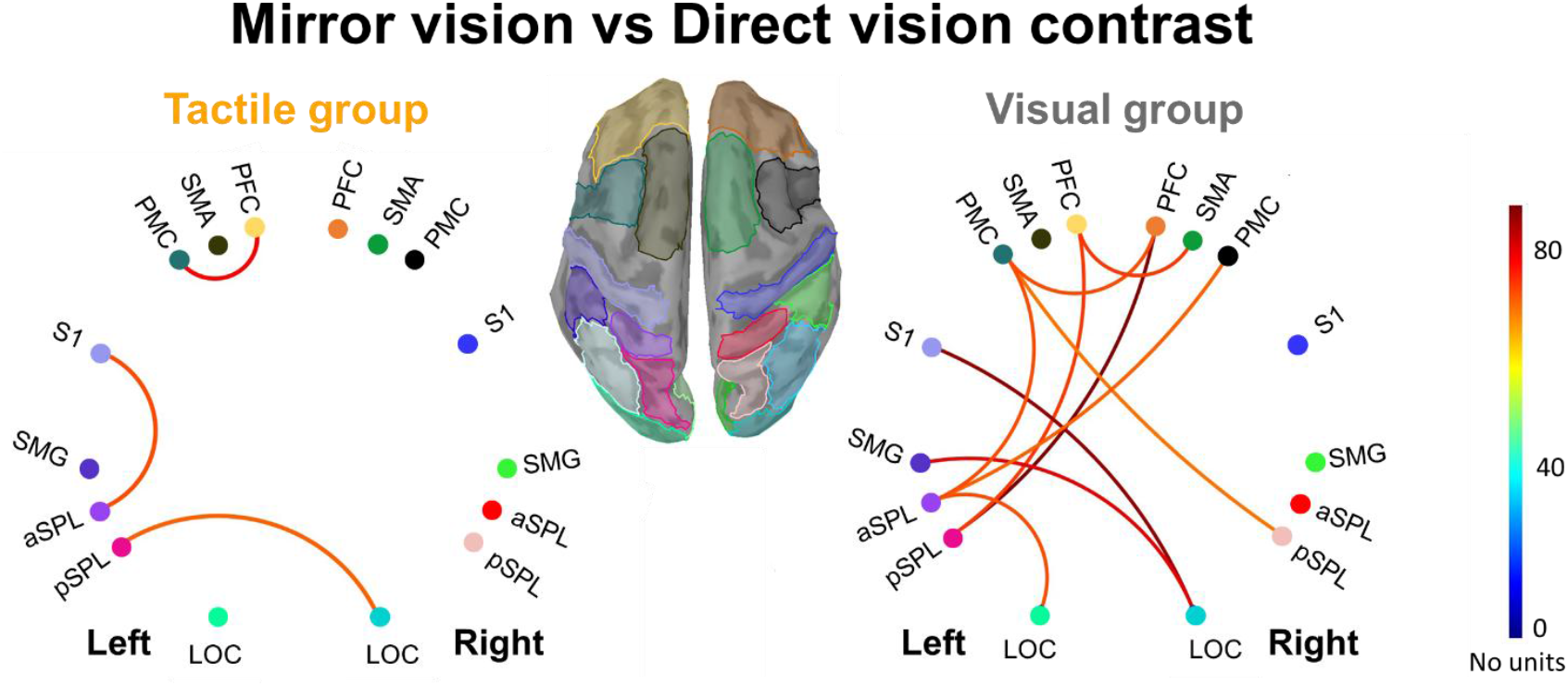
Spatial tactile cues influence cortical network recruitment during mirror tracing. Statistical functional connectivity among sixteen predefined ROIs during the tracing phase, displaying significant couplings resulting from the contrast between the Mirror and Direct vision conditions for the Tactile group (left) and the Visual group (right). aSPL: anterior superior parietal lobule, LOC: lateral occipital cortex, PFC: prefrontal cortex, PMC: premotor cortex, pSPL: posterior superior parietal lobule, S1: somatosensory cortex, SMA: supplementary motor area, SMG: supramarginal gyrus.

Statistical connectivity analyses revealed that the Tactile group engaged a selective network during mirror-reversed tracing. This network exhibited strengthened functional coupling between regions involved in somatosensory (S1–aSPL) and visual (LOC–pSPL) processing, as well as between regions operating at different levels of the motor control hierarchy (PFC–PMC) (Fig. 5, left panel). By contrast, in the Visual group, sensory conflict was associated with a more distributed and bilateral connectivity pattern, marked by pronounced cross-hemispheric interactions and strong involvement of frontal regions (Fig. 5, right panel).

## Discussion

When vision and touch convey conflicting spatial information during movement, somatosensory input is typically downweighted as unreliable^11,12,29^. Our findings demonstrate that introducing spatially structured tactile signals fundamentally reshapes this sensory weighting. In their presence, participants no longer exhibited the characteristic deterioration in motor control induced by mirror-reversed vision; instead performance remained comparable to that observed under direct vision. At the neural level, the conflict-related suppression of somatosensory inputs in S1 was reversed, with a significant increase in S1 activity. Together, the preserved motor performance and enhanced somatosensory activity indicate that explicitly structured tactile information transforms otherwise down-weighted or unreliable somatosensory signals into a potent resource for guiding movement under sensory conflict.

Recent evidence indicates that touch conveys directional information that can guide sliding finger movements^11,30–32^, challenging the traditional view that tactile input primarily serves perceptual rather than motor functions^33^. Notably, this motor-relevant contribution persists even when the hand is visible^11^, a context in which vision is typically assumed to dominate^34^. However, because somatosensory processing impairs movement performance under visuo-somatosensory conflict, the effect of tactile signals on motor control may become detrimental when tactile and visual information are incongruent^11,12,29^. In this sensory-conflict context, and in line with sensory-weighting theories ^35–40^, the reduction in somatosensory activity reported by Vlachou et al. (2025), and replicated here when tactile input lacked spatial information, suggests that motion-related tactile cues were treated as low-reliability, task-irrelevant signals and were effectively down-weighted during tracing movements.

This raises a critical question: are the disruptive effects of tactile processing an inherent consequence of visuo-somatosensory conflict, or do they depend on the informational content of the tactile signal? To address this, we adapted the mirror-tracing paradigm^11^ to introduce tactile spatial structure through discrete texture changes aligned with the polygon outline traced by participants with their index finger. When tactile feedback contained this explicit spatial information (Tactile group), tracing accuracy and smoothness were preserved under mirror-reversed vision matching performance observed under direct vision. By contrast, participants receiving tactile input that conveyed motion but lacked task-relevant spatial cues (Visual group) showed a marked deterioration in performance. Notably, the Tactile group maintained near-optimal tracing performance from the first block of trials, with stable movement accuracy and smoothness throughout the session. The Visual group, in contrast, exhibited gradual improvement in accuracy, as typically observed with repeated exposure to sensory conflict^12,41^. Yet, by the end of the session, their movement accuracy and smoothness remained significantly lower than those of the Tactile group. Overall, these findings suggest that task-relevant tactile spatial structure can rapidly resolve visuo-somatosensory conflict, providing an effective control signal for hand movements, yielding performance that surpasses what can be achieved through gradual sensorimotor adaptation alone.

The facilitative effect of spatial tactile cues in the mirror-tracing task is unlikely to be explained by mechanical constraints imposed by the raised dots marking the polygon outline. Their height (0.6 mm), comparable to standard Braille dots designed for perceptual discrimination rather than movement guidance, is too small to physically channel or deflect the fingertip. Instead, the finger pad simply deforms and slides over these dots, rather than being mechanically constrained by them (see Corniani et al. 2025, for visualization of skin deformation induced by fine surface textures)^42^. Therefore, the enhanced performance of the Tactile group reflects the informational content of the spatial tactile cues rather than physical restrictions on finger motion.

Source-space EEG analyses further indicate that, in presence of explicit spatial tactile cues, the visuo-somatosensory conflict modulated multiple stages of tactile processing. Notably, activity increased in the primary somatosensory cortex (S1), the principal cortical recipient of thalamic tactile input^1^, consistent with an early-stage enhancement of spatially informative tactile signals. In parallel, activity increased in the right posterior parietal cortex (PPC), a region critically involved in visuospatial processing, spatial attention, multisensory integration^9,43,44^, and in the transformation of tactile information into external reference frames^8,9,45^. These results support a framework in which S1 functions not merely as a passive relay but dynamically upregulates the effective neuronal weighting of behaviourally relevant somatosensory inputs when spatial remapping is required, as under visuo-somatosensory conflict. This early amplification may have facilitated the integration of tactile signals within PPC and their transformation into external coordinates. At the network level, S1 upregulation may also contribute to the enhanced functional connectivity observed in the Tactile group between S1 and the left aSPL, a region implicated in higher-level processing of environment-related tactile information^45^.

The Tactile group also exhibited strengthened functional coupling between the lateral occipital cortex (LOC) and posterior superior parietal lobule (pSPL), as well as between prefrontal cortex (PFC) and premotor cortex (PMC), under sensory conflict. Enhanced LOC-pSPL coupling reflects increased information exchange between regions involved in visuo-haptic integration (LOC)^46^ and adaptive hand motor control (pSPL)^47,48^. Although the present non-directional functional connectivity analyses do not reveal the direction of information flow, the strengthened PFC-PMC coupling may reflect top-down control from the PFC, a network implicated in the deliberate selection, updating, and monitoring of action plans^49,50^. Together, these neuronal modulations suggest that explicit spatial tactile cues enhance both the extrinsic encoding of somatosensory input and hierarchical sensorimotor coordination, yielding a network configuration that is both selective and efficient for guiding movement under conditions of sensory conflict.

In marked contrast, participants in the Visual group, who lacked spatially informative tactile cues under mirror-reversed tracing, exhibited reduced activity in both S1 and aSPL, replicating our previous findings^11^. In that earlier study, somatosensory gating occurred when participants traced the shape with the bare index finger, but not when a finger splint eliminated tactile information about finger-surface motion. Because both the current and previous manipulations selectively altered tactile inputs, the observed gating in S1 most likely reflects a stronger suppression of tactile, rather than proprioceptive signals. Despite this gating, persistently higher tracing errors and increased movement jerk under mirror vision indicate that somatosensory suppression only partially mitigated the sensory conflict in the Visual group, likely preventing an even greater deterioration of motor performance.

The absence of task-relevant spatial tactile information under sensory conflict was also associated with decreased activity in the left pSPL, a region critical for constructing spatial body representations, integrating multisensory information for action^9,27^ and expressing movement plans in visually-defined reference frames^51–53^. The functional significance of this reduction remains open to interpretation, with at least two non-exclusive possibilities. First, effective multisensory integration within pSPL may require compatibility between somatosensory feedback and the spatial structure of visual information, such that pSPL involvement is limited when this compatibility is violated. Alternatively, the reduced activity may reflect active attenuation of pSPL engagement to prevent conflicting sensory inputs from contaminating state estimation. From this perspective, such down-regulation may serve as an adaptive mechanism, reducing the computational costs of resolving conflict arising from incongruent sensory cues and allowing the motor system to rely more on stable sources of information, such as vision or internal predictions, to guide action. Notably, because activity in visual areas did not differ significantly between the Mirror and Direct vision conditions, the combined reduction in S1 and pSPL activity may have indirectly increased the relative weighting of visual feedback, contributing to a partial, though incomplete, reduction in the impact of sensory conflict on tracing performance.

Motor and prefrontal regions in the Visual group also exhibited reduced activation during sensory conflict, despite their established roles in movement planning, online monitoring, and computation of internal model predictions^54^. Given the increased demand for error detection and corrective control during visuo-somatosensory conflict, this reduction may appear counterintuitive. However, recent evidence suggests that high sensory uncertainty can suppress frontal and premotor activity to prevent overcorrection and increased movement variability^55^. Within this framework, the observed reduction in motor-area activity may reflect down-regulation of corticospinal drive when prediction errors become unreliable—a mechanism conceptually analogous to the reduced motor cortex excitability seen in freezing behaviours, known to prevent the generation of unstable motor commands^56^.

At the network level, a markedly different functional connectivity pattern emerged when explicit spatial tactile cues were absent during mirror tracing. In contrast to the relatively constrained network observed when relevant tactile spatial cues were available, the Visual group exhibited a largely distributed, cross-hemispheric connectivity pattern spanning frontal, parietal and occipital cortices. This shift toward large-scale network engagement suggests that high sensory uncertainty promotes distributed rather than localized neural communication^57,58^. The connectivity profile observed in the Visual group likely supports interactions necessary for sensory-guided action selection and updating, as well as internal exploration of competing alternatives^59–61^.

Compared to the Direct vision condition, Mirror vision condition was also associated with stronger functional connectivity between LOC and regions central to somatosensory processing (S1, SMG and aSPL), indicating enhanced cross-modal interactions. Given the established involvement of the LOC in hand-related visual representations^62,63^, this increased coupling may reflect a partial recoding of unreliable, somatosensory-based egocentric finger representations into more reliable, visually defined spatial coordinates under sensory conflict.

Overall, our results support the view that multisensory integration is inherently flexible: rather than operating through fixed, mandatory fusion, the contribution of each modality is actively and continuously adjusted according to task demands and contextual relevance^21,72, 73,74^. This integration may be mediated by top-down attentional mechanisms that are recruited in response to the estimated reliability and task relevance of sensory inputs, selectively facilitating informative signals and gating unreliable ones^65,66^. During motor execution, attentional focus can be directed internally toward one’s own movements or externally toward environmental cues, with external focus consistently improving movement accuracy, coordination and efficiency^67,68^. Although attention was not explicitly controlled in our study, the concurrent availability of explicit spatial tactile cues and spatiotopically coherent visual information during mirror-reversed tracing may have shifted participants’ focus toward task-relevant external features, stabilizing tracing performance. Attentional allocation to a specific modality is known to increase its perceptual reliability^69,70^, by amplifying context-relevant neural responses in both early sensory^71,72^ and high-order associative^73–75^ areas. Accordingly, the cortical modulations observed here may reflect differential weighting driven by attentional allocation, depending on the informational content of the tactile cues.

By demonstrating that explicit spatial tactile cues can transform otherwise unreliable somatosensory signals into task-relevant inputs, our findings provide a conceptual framework for further clinical applications. Future studies could investigate whether augmenting spatiotopic or spatially structured tactile feedback supports motor performance in neurological populations characterised by impaired somatosensory processing or disrupted internally generated movements, including stroke^76^, hand dystonia^77^, and Parkinson’s disease^56^. Leveraging spatial tactile cues may therefore offer a practical strategy for enhancing motor rehabilitation by promoting more reliable multisensory integration and more precise motor control.

## Methods

### Participants

Thirty-seven adults participated in the study (16 men, 21 women; mean age ± SD = 24.7 ± 3.1 years), which lasted approximately 2 hours. All participants were right-handed, as confirmed by the Edinburgh Handedness Inventory^78^ (mean laterality score ± SD = 81.1 ± 22), and reported normal or corrected-to-normal vision, with no history of sensory or motor impairments. The sample size was chosen based on previous studies examining hand motor control under mirror-reversed visual feedback^11,13^. Participants were randomly assigned to one of two experimental groups (Tactile group: 8 men, 11 women; mean age ± SD = 24.5 ± 2.74 years, and Visual group: 8 men, 10 women; mean age ± SD = 24.9 ± 3.5 years). Following completion of the experiment, they received monetary compensation for their participation. The study was approved by the CERSTAPS Ethics Committee (no. IRB00012476-2021-09-12-140) and conducted in accordance with the principles of the Declaration of Helsinki. Written informed consent was obtained from all participants prior to the experiment.

### Experimental set up

Participants sat comfortably at a table in a dimly lit room, with a light source positioned above the workspace (mean ambient temperature: 24.7 ± 2.4 °C, relative humidity: 44.9 ± 9.1% (RS PRO RS-172TK temperature-humidity meter, China)). Given that skin hydration affects skin mechanics and, consequently, the quality of tactile stimulation during finger-surface interactions^79^, we assessed the hydrolipid film of each participant’s right index fingertip using a skin moisture and oil content analyser (Hurrise, China). Measurements were collected at three time points: at the start of the experimental session, between the Mirror and Direct vision conditions, and at the end of the session (see below for the temporal organization of the experimental session). For each participant and each time point, the three measurements were averaged to obtain a single estimate. All participants exhibited similar skin hydration and oil levels within the normal physiological range^80^ (Supplementary Material: Fig. S1; Tables S16 & S17).

A circular mirror (Comair Cabinet Executive, diameter 280 mm) was positioned 45° to the left of the participants’ midline. Placed in front of them was a force plate (150 × 150 mm; AMTI HE6×6, A-Tech Instruments Ltd., Toronto, Canada) on which a 3D-printed surface (Sigma D25 printer, BCN3D) made of biopolymer thermoplastic (polylactic acid, PLA; 195 × 195 × 2 mm) was screwed. The surface featured the white outline of an irregular polygon against a black background, which participants traced using the pulp of their index finger. The polygon consisted of 17 segments of varying lengths and had a total perimeter of 652 mm (see inset in Fig. 1a). Each segment was composed of three rows of white Braille-like dots (dot diameter: 1.5 mm; height: 0.6 mm; inter-dot spacing: 2.5 mm). Such Braille-like textures have been shown to effectively activate mechanoreceptors involved in tactile perception^81^.

Participants were assigned to one of two experimental groups that differed in the texture properties of the surface used for tracing. In the Tactile group, the contour of the white polygon was composed of raised Braille-like dots and contrasted with the smooth black background (Fig. 1b). This configuration provided both visual and tactile cues, allowing participants to haptically and visually discern whether their finger was on or off the outline. In the Visual group, the entire surface, including the area surrounding the polygon, was uniformly covered with the same Braille-like texture (Fig. 1c). In this case, the only feature distinguishing the outline was its colour (white outline on black background).

### Task paradigm

Before the experimental session, participants washed and thoroughly dried their hands to minimize variability in skin hydration levels. Following this preparation, and to establish a baseline, the session began with a control condition, during which participants followed the outline of the polygon as accurately as possible while directly viewing their hand and the polygon (Direct vision condition). At the beginning of the first trial, participants placed their index finger on the point of the polygon indicated by a yellow arrow in Fig. 1a. During the first 3 s, they adjusted the normal force applied to the surface to remain within the 0.2 - 0.4 N range, consistent with natural tactile exploration^82^. Force levels were monitored in real time on the experimenter’s display and corrective feedback was provided when necessary. Offline analyses confirmed that participants in both groups maintained stable normal force during tracing, with no substantial differences in force levels between groups (Supplementary Fig. S2 & Table S18), thereby avoiding large variations known to modulate somatosensory cortex activity ^83–85^. After normal force stabilization, an auditory signal (a beep) marked the start of a 7 s static phase, during which participants remained motionless to establish a baseline for the EEG analyses. A second beep cued them to begin tracing the polygon outline, until a final beep, occurring 30 s later, indicated the end of the trial. For each subsequent trial, participants resumed tracing from the point at which they had stopped in the previous trial.

After completing the Direct vision condition, participants performed the same tracing task while viewing their hand and the polygon through the mirror (Mirror vision condition). Direct vision of both the hand and the tracing surface was blocked with a black shield (Fig. 1a). This mirror setup introduces a discrepancy between visual and somatosensory inputs, typically resulting in less accurate and jerkier movements^11,12,86,87^. Participants were required to maintain their index fingertip on the polygon outline throughout the task. In case of deviation, they had to return to the point of deviation before continuing the tracing.

In both visual conditions, participants were instructed to remain relaxed, minimize eye movements, and prioritize finger and wrist movements over arm movements. In line with prior EEG studies involving sensory conflict^11–13,18^, they were asked to trace the polygon slowly (∼10 mm/s), a speed too low to complete the entire shape in a single trial. This slow speed was chosen to prevent somatosensory attenuation during rapid movements^88^, minimise contamination of EEG signals by muscle or ocular artefacts, and promote comparable tracing speeds across the Direct Vision and Mirror vision conditions. Before data collection, the experimenter demonstrated the target tracing speed and monitored compliance throughout the session.

Participants of both groups completed 20 trials per condition (Mirror vision and Direct vision). Because our EEG analyses relied on averaging across trials within each condition, the protocol was designed to limit adaptation to the novel visuo-somatosensory environment reported in previous studies^12,41^, thereby ensuring that condition-averaged data remained representative. As described above, participants progressed along the polygon by resuming each trial from the point reached at the end of the previous one rather than returning to the initial starting point, a procedure implemented to avoid repeatedly tracing the same segments and thus reduce practice-related learning. After every five trials, participants took a brief (∼20 s) pause during which they moved and directly viewed their hand. The irregular polygon, with its numerous directional changes, increased task complexity^20^, thereby further limiting adaptation to the sensory conflict across trials.

### Data acquisition and analysis

#### Task performance

Participant movements were recorded using a GoPro Hero Session camera (Full HD, 1920 × 1080 pixels; 60 fps) positioned 290 mm above and parallel to the surface, capturing the full trajectory of the index fingertip during tracing. Video data were analysed with Kinovea, an open-source motion-tracking and annotation software. A small cross marked on the fingernail (see below) served as the visual marker tracked in Kinovea. Spatial calibration was verified by confirming that the polygon perimeter extracted from the video matched its actual physical length, ensuring accurate spatial scaling. For each trial, movement data were analysed from the onset of movement following the auditory “start tracing” signal until the offset of movement at the “stop tracing” signal.

Reaction forces and moments exerted by the index finger on the surface were recorded at 256 Hz using an AMTI HE6 × 6 force plate, on which the 2 mm-thick surface was screwed. This compact device provides high resolution (12-bit) measurements of low-magnitude forces (up to 2.2 N in Fx and Fy, and 4.5 N in Fz), making it well suited for capturing the small forces applied by the fingertip during tracing. Force data were digitized using a Keithley 12-bit A/D converter (AD-win pro, Keithley Instruments, Cleveland, OH) and processed offline in MATLAB 7.0 (The MathWorks, Natick, MA).

Tracing accuracy was assessed using the error index, defined as the ratio between the total distance traced by the fingertip and the actual length of the corresponding portion of the polygon outline traced during the trial. Movement smoothness was quantified using the logarithmic dimensionless jerk (LDLJ) index, derived from the rate of change of the resultant fingertip force in the horizontal (XY) plane. This metric provides an objective measure of the stability and abruptness of force transitions during the task, capturing both sudden and intermittent fluctuations. While traditionally employed in the assessment of sensorimotor impairments^76,89–91^, recent studies have applied jerk-based metrics to characterize motor performance in behavioural tasks^11,92,93^.

For each trial, LDLJ was calculated as:

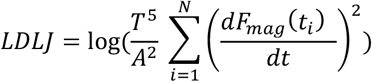

where 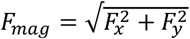 is the magnitude of the resultant force in the XY plane, *T* is the trial duration, *A* is the total force amplitude (defined as the difference between the maximum and minimum *Fmag* within each trial, and *dFmag/dt* is the first derivative of the resultant force over time. Because reaction times to the “start tracing” signal varied across participants, LDLJ was particularly appropriate for this task, as it normalizes for both the magnitude and duration of force changes, enabling consistent comparisons of movement smoothness across trials, participants and experimental conditions.

#### Electroencephalography (EEG)

Continuous EEG activity was recorded from 64 Ag/AgCl electrodes with integrated preamplifiers, mounted on an elastic cap (BioSemi ActiveTwo system, Amsterdam, The Netherlands). Five participants were excluded from the EEG analysis due to technical issues or excessive artefacts. The final EEG dataset included 32 participants: Tactile group (6 men, 10 women; mean age ± SD = 24.07 ± 2.7 years) and Visual group (8 men, 10 women; mean age ± SD = 24.8 ± 3.6 years).

In the BioSemi system, the conventional ground electrode is replaced by a Common Mode Sense (CMS) active electrode and a Driven Right Leg (DRL) passive electrode, which together form a feedback loop to stabilize the common-mode voltage. Signals were digitized at 1024 Hz with a 24-bit resolution and recorded using ActiView software. Four additional Ag/AgCl electrodes were placed near the outer canthi, and above and below the left eye to record electrooculographic (EOG) activity, which was subsequently used to monitor and correct ocular artefacts contaminating the EEG signals.

Pre-processing was performed in BrainVision Analyzer 2 (Brain Products, Gilching, Germany). The EEG data were down-sampled to 200 Hz and band-pass filtered (0.5–90 Hz), with a 50 Hz notch filter applied to remove line noise. Channels with continuous poor signal quality were interpolated. Signals were re-referenced to the average of all scalp electrodes. Ocular and blink artefacts were corrected using independent component analysis (ICA), and the cleaned data were subsequently exported to Brainstorm^94^ for source-space analysis.

In Brainstorm, EEG signals were segmented into epochs time-locked to the “start tracing” tone for each experimental condition. Each epoch included a 2 s static pre-tracing baseline and a 30 s tracing period (see Fig. 6a). Epochs were baseline-corrected using the static pre-tracing period. All trials and channels were visually inspected for artefacts; bad channels and noisy trials were discarded. On average, 17.3 ± 2.6 (SD) epochs of 32 s per participant were retained for analysis (Direct condition: 17.2 ± 2.9; Mirror condition: 17.4 ± 2.3).

**Fig. 6.**
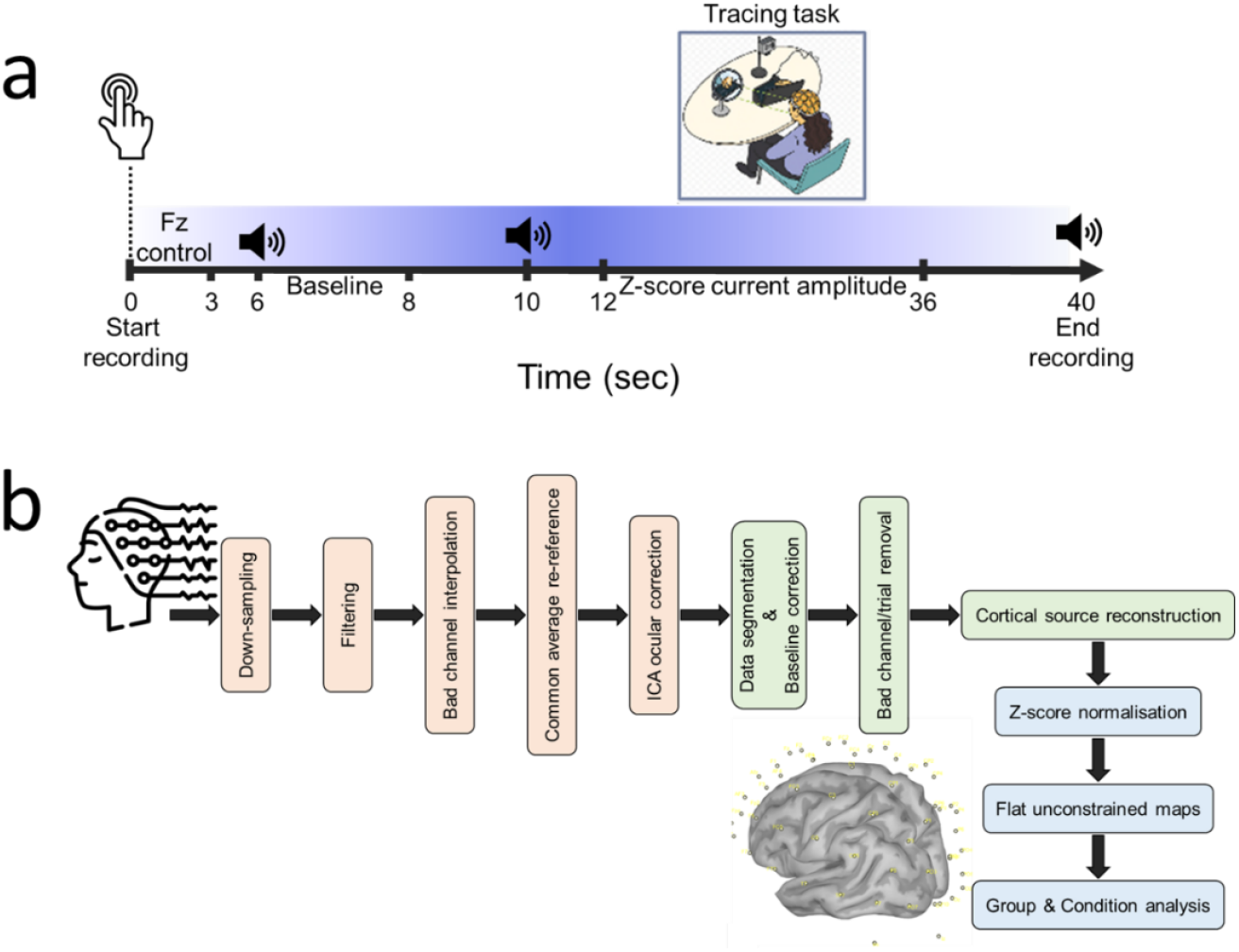
**a Experimental timeline of each trial**. The EEG signals recorded during the tracing task were normalized using z-score transformation of the current computed in the 2s time window (Baseline) of the Static phase. The z-scored current amplitude was then averaged over the 12-36s time window. **b Schematic overview of the EEG signal processing and analysis pipeline**. Steps shown in orange were performed using BrainVision Analyzer, steps in green were carried out in Brainstorm, and steps in blue represent analyses conducted in source space.

Cortical source reconstruction was performed using the Colin27 MRI template (Montreal Neurological Institute) and the BioSemi 64-channel 10–10 electrode layout. The forward model was computed with a three-shell spherical head model (scalp, outer skull, inner skull; 15,002 vertices), which accounts for conductivity differences between tissues^95^. The inverse solution was estimated using minimum norm estimation (MNE) with unconstrained dipole orientations, followed by dynamic statistical parametric mapping (dSPM) normalization.

In source space, current amplitude was normalized using a z-score transformation relative to the mean activity during the static baseline (6–8 s in Fig. 6a). The three components (x, y, z) at each source location were combined using Principal Component Analysis (PCA) to generate flat maps. Source analyses were performed over the entire cortical surface and within a selected set of regions of interest (ROIs). Specifically, we manually defined 16 regions of interest (ROIs), corresponding to eight bilateral cortical areas known to contribute to visually guided tracing, visuo-somatosensory integration, and sensory conflict resolution^25–28^: the anterior superior parietal lobule (aSPL), lateral occipital cortex (LOC), prefrontal cortex (PFC), premotor cortex (PMC), posterior superior parietal lobule (pSPL), somatosensory cortex (S1), supplementary motor area (SMA), and supramarginal gyrus (SMG). For each ROI, source-level activity was quantified as the mean absolute current amplitude during the 2-26 s interval of the 30-second tracing phase (Fig. 6b for a detailed overview of the EEG analysis pipeline). This window was deliberately chosen to exclude transient responses to the auditory “start” and “stop” tones marking the tracing phase and to accommodate individual variability in tracing onset times. Tracing performance under mirror-reversed visual feedback is not uniform throughout the task, as it can become more challenging at corners or specific orientations^13,96^. Accordingly, averaging current amplitude across the 2-26 s tracing interval provides a conservative estimate of the effects of sensory conflict.

### Statistical analysis

Statistical analysis was conducted using JASP (Version 0.95.4, University of Amsterdam) and STATISTICA 8.0 (StatSoft, Inc., USA) with detailed outputs provided in the Supplementary Material (Table S1 – S18). Data normality was assessed using the Shapiro–Wilk test, and homogeneity of variances was confirmed with Levene’s test. When normality assumptions were violated, data were log-transformed to meet parametric test assumptions. If these assumptions remained unmet after transformation, non-parametric alternatives were applied, including the Wilcoxon signed-rank test, Friedman test, Mann–Whitney U test, Conover’s post hoc test, as appropriate for the experimental design.

Behavioural variables were analysed using 2 (Group: Tactile, Visual) × 2 (Condition: Direct vision, Mirror vision) mixed ANOVAs, with Condition as the within-subject factor and Group as the between-subject factor. Significant main or interaction effects were further explored with post hoc pairwise comparisons using Fisher’s least significant difference (LSD) test, with the alpha level set at .05. Effect sizes are reported as partial eta squared (η^2^_p_) for ANOVAs, and as Cohen’s d or Rank-Biserial Correlation (r_rb_) for pairwise comparisons, depending on the statistical test. The significance threshold was set at p < .05 for all analyses.

Within-subject EEG analyses contrasted the absolute source-estimated current amplitude between the Mirror and Direct vision conditions using a two-tailed paired permutation Wilcoxon signed-rank test (1000 randomizations; significance threshold p < 0.05, uncorrected). For between-group comparisons, average current amplitudes were extracted from predefined regions of interest (ROIs) and submitted to 2 (Group: Tactile, Visual) × 2 (Condition: Direct vision, Mirror vision) mixed ANOVAs, with Condition as the within-subject factor. Significant interaction effects were followed up with Holm-corrected post hoc comparisons.

Data visualization was performed in R (RStudio 2025.05.1+513) using the ggplot2 and vioplot packages.

## Data availability

The data supporting the findings of this study are available from the corresponding author upon reasonable request.

## Acknowledgements

The project leading to this publication has received funding from the French government under the “France 2030” investment plan managed by the French National Research Agency (reference: ANR-16-CONV000X / ANR-17-EURE-0029), from Excellence Initiative of AixMarseille University - A*MIDEX (AMX-19-IET-004) and the project COMTACT (ANR 2020-CE28-0010-03), funded by the French “Agence Nationale de la Recherche” (ANR). We thank Dany Paleressompoulle for the fabrication and 3-D printing of the surfaces, Franck Buloup for developing the Docometre software used for data acquisition, and we are grateful to all participants for their involvement in the study.

## Author contributions

B.J., V.M-E., and M.L. conceived and designed the study. V.M-E. and L.E. collected and analysed the data. V.M-E. drafted the manuscript. V.M-E., B.J., M.L. reviewed and edited the manuscript.

## Declarations Competing Interests

The authors declare no competing interests.

## Materials & Correspondence

Correspondence and requests for materials should be addressed to V.M-E.

## Acknowledgments

We are grateful to Emerson College for administering the survey. Specifically, Patrick Fox, Matt Taglia, Camille Mumford, and Spencer Kimball. We thank the following persons for their methodological advisement: Stephen Immerwahr and Sungwoo Lim of the NYC Health Department, Dr. Lauren Russell of the Federal Reserve Board, Dr. Raffi Garcia of Rensselaer Polytechnic Institute, Dr. Gwendolyn Wright of Duke / Cook Center, Dr. Sara Chaganti and Dr. Mike Evangelist of the Federal Reserve Bank of Boston, Dr. Laura Sullivan of the New Jersey Institute for Social Justice, Dr. Suparna Bhaskaran and Elaine Chang of the New School Institute for Race, Power, and Policy, and Dr. Chris Wimer and Schuyler Ross of Columbia University.

## Notes

### Competing Interest Statement

The authors have declared no competing interest.

