## Supplementary material for "Making sense of touch: spatial tactile cues promote efficient visuo–somatosensory integration and restore motor accuracy under sensory conflict"

### List of supplementary material

Page

*Table S1. **Error index**: Within & between subjects effects.*

| Effect | Sum of squares | df | Mean Square | F | p | $\eta^2p$ |
|---|---|---|---|---|---|---|
| **Intercept** | 106.92 | 1 | 106.92 | 4760.46 | 0.00 | 0.99 |
| **Group** | 0.31 | 1 | 0.31 | 13.99 | <.001*** | 0.29 |
| **Error** | 0.79 | 35 | 0.02 | | | |
| **Condition** | 0.63 | 1 | 0.63 | 24.00 | <.001*** | 0.41 |
| **Condition * Group** | 0.24 | 1 | 0.24 | 9.2 | 0.005** | 0.21 |
| **Error** | 0.9 | 35 | 0.03 | | | |

*Table S2. **Error index**: Hoc comparisons – Group ✻ Condition.*

| Cell number | Group | Condition | {1} 1.1 | {2} 1.17 | {3} 1.12 | {4} 1.42 |
|---|---|---|---|---|---|---|
| **1** | Tactile | Direct vision | | 0.19 | 0.75 | <.001*** |
| **2** | Tactile | Mirror vision | 0.19 | | 0.3 | <.001*** |
| **3** | Visual | Direct vision | 0.75 | 0.3 | | <.001*** |
| **4** | Visual | Mirror vision | <.001*** | <.001*** | <.001*** | |

** p < .01, *** p < .001.

**Tables S1-S2** Results of the 2 (Group: Tactile, Visual) × 2 (Condition: Direct vision, Mirror vision) mixed ANOVA, with repeated measures on the factor Condition on the error (distance-to-segment) index. The analysis revealed a significant Group × Condition interaction. Post hoc pairwise comparisons using Fisher's LSD indicated that error index remained stable across vision conditions in the Tactile group, whereas it increased significantly under Mirror vision in the Visual group. Analyses were conducted on the logarithmic values of the data.

Table S3. **Jerk:** Within & between subjects effects.

| Effect | Sum of squares | df | Mean Square | F | p | $\eta^2p$ |
|---|---|---|---|---|---|---|
| **Intercept** | 52985.67 | 1 | 52985.67 | 89988.33 | 0.00 | 0.99 |
| **Group** | 0.54 | 1 | 0.54 | 0.92 | 0.34 | 0.03 |
| **Error** | 20.61 | 35 | 0.59 | | | |
| **Condition** | 0.15 | 1 | 0.15 | 5.55 | 0.02* | 0.14 |
| **Condition * Group** | 0.39 | 1 | 0.39 | 13.89 | <.001*** | 0.28 |
| **Error** | 0.98 | 35 | 0.03 | | | |

*Table S4. **Jerk:** Hoc comparisons - Group ✲ Condition.*

| Cell number | Group | Condition | {1} 26.71 | {2} 26.67 | {3} 26.74 | {4} 26.97 |
|---|---|---|---|---|---|---|
| **1** | Tactile | Direct vision | | 0.33 | 0.89 | 0.16 |
| **2** | Tactile | Mirror vision | 0.33 | | 0.67 | 0.09 |
| **3** | Visual | Direct vision | 0.89 | 0.67 | | <.001*** |
| **4** | Visual | Mirror vision | 0.16 | 0.09 | <.001*** | |

* p < .05, *** p < .001.

**Tables S3-S4** Summary of the 2 (Group: Tactile, Visual) × 2 (Condition: Direct vision, Mirror vision) mixed ANOVA, with repeated measures on the factor Condition on the jerk index (log dimensionless jerk-LDLJ). The analysis revealed a significant Group × Condition interaction. Post hoc pairwise comparisons using Fisher's LSD showed that jerk remained stable across vision conditions in the Tactile group, whereas a significant increase was observed in the Visual group during the Mirror vision condition.

*Table S5. **Error index ratio // Tactile group:** Friedman test.*

| Factor | $X^2_F$ | DF | P | KENDALL'S W |
|---|---|---|---|---|
| **Block of trials** | 0.467 | 3 | .926 | 0.009 |

*Table S6. **Error index ratio // Visual group:** Friedman test.*

| Factor | $X^2_F$ | DF | P | KENDALL'S W |
|---|---|---|---|---|
| **Block of trials** | 14.47 | 3 | .002** | 0.268 |

*Table S7. **Error index ratio // Visual group:** Conover's Post hoc comparisons – block of trials.*

| Block | | T-Stat | df | $W_i$ | $W_j$ | $r_{rb}$ | $p_{holm}$ |
|---|---|---|---|---|---|---|---|
| **1** | 2 | 1.466 | 51 | 58.00 | 48.00 | 0.637 | .297 |
| | 3 | 1.906 | 51 | 58.00 | 45.00 | 0.556 | .187 |
| | 4 | 4.252 | 51 | 58.00 | 29.00 | 0.801 | < .001*** |
| **2** | 3 | 0.440 | 51 | 48.00 | 45.00 | 0.251 | .662 |
| | 4 | 2.786 | 51 | 48.00 | 29.00 | 0.708 | .037* |
| **3** | 4 | 2.346 | 51 | 45.00 | 29.00 | 0.474 | .092 |

* p < .05, ** p < .01, *** p < .001.

*Note.* Rank-biserial correlation based on individual signed-rank test.

**Tables S5-S7** Summary of two separate non-parametric Friedman tests (within-group repeated-measures analyses across Blocks 1–4; each block comprising five consecutive trials) assessing adaptation in the error index ratio (Mirror/Direct vision conditions) for the Tactile and Visual groups. The Tactile group showed no significant main effect of Block, indicating stable performance across the session in presence of explicit spatial tactile cues. In contrast, the Visual group exhibited a significant Block effect. Conover's post hoc tests revealed that error indices in Blocks 1 and 2 were significantly higher than in Block 4, indicating improved accuracy in the last trials of the session in the Visual group.

*Table S8.* ***Error index ratio:*** *Independent samples t-test.*

| Block | U | P | Rank-Biserial Correlation | SE Rank-Biserial Correlation |
|---|---|---|---|---|
| **1** | 79.00 | .004** | 0.538 | 0.190 |
| **2** | 53.00 | < .001*** | 0.690 | 0.190 |
| **3** | 65.00 | < .001*** | 0.620 | 0.190 |
| **4** | 94.00 | .031* | 0.420 | 0.193 |

* p < .05, ** p < .01, *** p < .001

Note. For the Mann-Whitney test, effect size is given by the rank biserial correlation.

Note. Mann-Whitney U test.

**Table S8** Summary of four independent non-parametric Mann–Whitney tests comparing the median error index ratio (Mirror/Direct vision conditions) between the Tactile and Visual groups for each of the four trial blocks. In each block, the Visual group exhibited a significantly higher error index ratio than the Tactile group, indicating consistently less accurate tracing performance throughout the session in the absence of explicit spatial tactile cues.

*Table S9.* ***Jerk:*** *Within & between subjects effects.*

| Effect | Sum of squares | df | Mean Square | F | p | $\eta^2p$ |
|---|---|---|---|---|---|---|
| **Intercept** | 148.93 | 1 | 148.93 | 479538.2 | 0.00 | 0.99 |
| **Group** | 0.004 | 1 | 0.004 | 13.4 | < .001*** | 0.28 |
| **Error** | 0.01 | 35 | 0.0003 | | | |
| **Block** | 0.0007 | 3 | 0.0002 | 2.3 | 0.09 | 0.06 |
| **Block * Group** | 0.0002 | 3 | 0.0001 | 0.5 | 0.65 | 0.02 |
| **Error** | 0.01 | 105 | 0.0001 | | | |

*** p < .001.

**Table S9** Results of the 2 (Group: Tactile, Visual) × 4 (Block: 1, 2, 3, 4) mixed ANOVA, with repeated measures on the factor Block assessing adaptation in the jerk ratio (Mirror/Direct vision conditions). The analysis revealed a significant main effect of Group, with the Visual group exhibiting higher jerk ratio values than the Tactile group. No significant main effect of Block or Block x Group interaction was found, indicating that movement smoothness remained stable across the session in both groups.

*Table S10. **Left postcentral gyrus (S1):** Within & between subjects effects.*

| Effect | Sum of squares | df | Mean Square | F | p | η²p |
|---|---|---|---|---|---|---|
| **Intercept** | 157.66 | 1 | 157.66 | 6275.55 | 0.00 | 0.99 |
| **Group** | 0.0003 | 1 | 0.0003 | 0.01 | 0.92 | 0.0004 |
| **Error** | 0.73 | 29 | 0.03 | | | |
| **Condition** | 0.01 | 1 | 0.02 | 1.41 | 0.24 | 0.05 |
| **Condition * Group** | 0.06 | 1 | 0.06 | 6.90 | 0.01* | 0.19 |
| **Error** | 0.24 | 29 | 0.01 | | | |

*Table S11. **Left postcentral gyrus (S1):** Hoc comparisons - Group ✳ Condition.*

| Cell number | Group | Condition | {1} 1.55 | {2} 1.64 | {3} 1.61 | {4} 1.58 |
|---|---|---|---|---|---|---|
| **1** | Tactile | Direct vision | | 0.01* | 0.17 | 0.50 |
| **2** | Tactile | Mirror vision | 0.01* | | 0.62 | 0.23 |
| **3** | Visual | Direct vision | 0.17 | 0.62 | | 0.32 |
| **4** | Visual | Mirror vision | 0.50 | 0.23 | 0.32 | |

* p < .05.

**Tables S10-S11** Results of a 2 (Group: Tactile, Visual) × 2 (Condition: Direct vision, Mirror vision) mixed ANOVA, with repeated measures on the factor Condition on the mean absolute source current amplitude in the left postcentral gyrus (S1) ROI. The analysis revealed a significant Group × Condition interaction. Post hoc pairwise comparisons (Fisher's LSD) indicated that current amplitude increased significantly from Direct to Mirror vision in the Tactile group, whereas no significant difference between conditions was observed in the Visual group.

*Table S12. **Left anterior superior parietal lobule (aSPL):** Within & between subjects effects.*

| Effect | Sum of squares | df | Mean Square | F | p | η²p |
|---|---|---|---|---|---|---|
| **Intercept** | 180.23 | 1 | 180.23 | 4859.80 | 0.00 | 0.99 |
| **Group** | 0.002 | 1 | 0.0017 | 0.05 | 0.83 | 0.002 |
| **Error** | 1.11 | 30 | 0.04 | | | |
| **Condition** | 0.04 | 1 | 0.04 | 1.38 | 0.25 | 0.04 |
| **Condition * Group** | 0.29 | 1 | 0.29 | 10.42 | 0.003** | 0.26 |
| **Error** | 0.84 | 30 | 0.03 | | | |

*Table S13.* ***Left anterior superior parietal lobule (aSPL):***
*Hoc comparisons - Group ✻ Condition.*

| Cell number | Group | Condition | {1} 1.64 | {2} 1.73 | {3} 1.77 | {4} 1.58 |
|---|---|---|---|---|---|---|
| **1** | Tactile | Direct vision | | 0.16 | 0.06 | 0.35 |
| **2** | Tactile | Mirror vision | 0.16 | | 0.55 | 0.03* |
| **3** | Visual | Direct vision | 0.06 | 0.55 | | 0.004** |
| **4** | Visual | Mirror vision | 0.35 | 0.03* | 0.004** | |

* p < .05, ** p < .01.

**Table S12–13** Results of a 2 (Group: Tactile, Visual) × 2 (Condition: Direct vision, Mirror vision) mixed ANOVA, with repeated measures on the factor Condition on the mean absolute source current amplitude in the left anterior superior parietal lobule (aSPL) ROI. A significant Group × Condition interaction was found. Post hoc pairwise comparisons (Fisher's LSD) revealed a significant decrease in aSPL activity for the Visual group in the Mirror vision condition relative to Direct vision, whereas no significant effect of mirror vision was observed in the Tactile group.

*Table S14.* ***Left posterior superior parietal lobule (pSPL):*** *Within & between subjects effects.*

| Effect | Sum of squares | df | Mean Square | F | p | $\eta^2p$ |
|---|---|---|---|---|---|---|
| **Intercept** | 176.13 | 1 | 176.13 | 4117.37 | 0.00 | 0.99 |
| **Group** | 0.00 | 1 | 0.00 | 0.00 | 0.99 | 0.00 |
| **Error** | 1.28 | 30 | 0.04 | | | |
| **Condition** | 0.12 | 1 | 0.12 | 5.22 | 0.03* | 0.15 |
| **Condition * Group** | 0.17 | 1 | 0.17 | 7.45 | 0.011* | 0.2 |
| **Error** | 0.70 | 30 | 0.02 | | | |

*Table S15.* ***Left posterior superior parietal lobule (pSPL):***
*Hoc comparisons - Group ✻ Condition.*

| Cell number | Group | Condition | {1} 1.65 | {2} 1.67 | {3} 1.75 | {4} 1.56 |
|---|---|---|---|---|---|---|
| **1** | Tactile | Direct vision | | 0.76 | 0.11 | 0.18 |
| **2** | Tactile | Mirror vision | 0.76 | | 0.18 | 0.11 |
| **3** | Visual | Direct vision | 0.11 | 0.18 | | 0.0013** |
| **4** | Visual | Mirror vision | 0.18 | 0.11 | 0.0013** | |

* p < .05, ** p < .01.

**Tables S14–15** Results of a 2 (Group: Tactile, Visual) × 2 (Condition: Direct vision, Mirror vision) mixed ANOVA, with repeated measures on the factor Condition on the mean absolute source current amplitude in the left posterior superior parietal lobule (pSPL) ROI. A significant Group × Condition interaction was found. Post hoc pairwise comparisons (Fisher’s LSD) revealed a significant decrease in pSPL activity for the Visual group in the Mirror vision condition relative to Direct vision, whereas no significant difference was observed in the Tactile group.

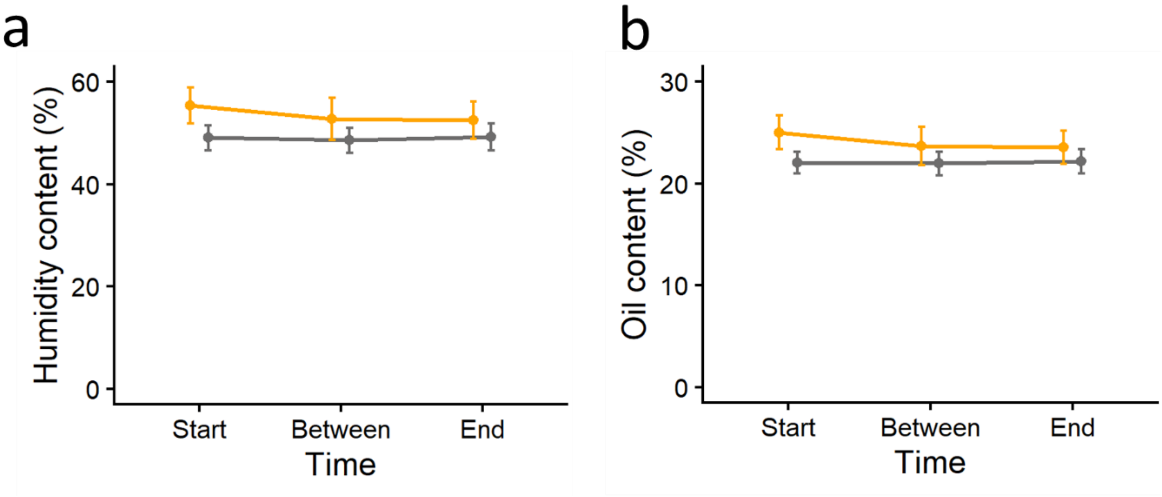

Fig. S1 | **Humidity and oil content of the index finger**. Line plots depict measurements of **(a)** humidity and **(b)** oil content for the Tactile (orange) and Visual (grey) groups at three time points: Start: before the experimental session; Between: between the Direct and Mirror vision conditions; End: at the end of the experimental session. Two 2 (Group: Tactile, Visual) × 3 (Time: Start, Between, End) mixed ANOVAs, with repeated measures on the factor Time, revealed no significant main effects of Group or Time and no Group × Time interactions, indicating comparable finger skin composition across groups and time points. Error bars denote ±95% confidence intervals of the mean.

*Table S16. **Humidity content:** Within & between subjects effects.*

| Effect | Sum of squares | df | Mean Square | F | p | $\eta^2p$ |
|---|---|---|---|---|---|---|
| **Intercept** | 234328.1 | 1 | 234328.1 | 2284.28 | 0.00 | 0.99 |
| **Group** | 374.3 | 1 | 374.3 | 3.65 | 0.07 | 0.12 |
| **Error** | 2872.3 | 28 | 102.6 | | | |
| **Time** | 41.8 | 2 | 20.9 | 2.03 | 0.14 | 0.07 |
| **Time * Group** | 32.0 | 2 | 16.0 | 1.56 | 0.22 | 0.05 |
| **Error** | 575.9 | 56 | 10.3 | | | |

**Table S16** Results of the 2 (Group: Tactile, Visual) × 3 (Time: Start, Between, After) mixed ANOVA, with repeated measures on the factor Time on mean finger skin humidity content measured at three time points: Start: before the experimental session; Between: between the Direct and Mirror vision conditions; End: at the end of the experimental session. The analysis revealed no significant main effect of Group or Time, and no significant Group × Time interaction, indicating that skin humidity did not differ between groups across the session. All analyses were performed on log-transformed humidity values.

*Table S17. **Oil content:** Within & between subjects effects.*

| Effect | Sum of squares | df | Mean Square | F | p | $\eta^2p$ |
|---|---|---|---|---|---|---|
| **Intercept** | 47512.68 | 1 | 47512.68 | 2237.84 | 0.00 | 0.99 |
| **Group** | 72.39 | 1 | 72.39 | 3.41 | 0.08 | 0.11 |
| **Error** | 594.48 | 28 | 21.23 | | | |
| **Time** | 8.39 | 2 | 4.20 | 1.94 | 0.15 | 0.06 |
| **Time * Group** | 9.20 | 2 | 4.60 | 2.12 | 0.13 | 0.07 |
| **Error** | 121.27 | 56 | 2.17 | | | |

**Table S17** Results of the 2 (Group: Tactile, Visual) × 3 (Time: Start, Between, After) mixed ANOVA, with repeated measures on the factor Time on mean finger skin oil content measured at three time points during the session. The analysis revealed no significant main effects and no Group × Time interaction, indicating that skin oil content did not differ between groups across the session. All analyses were performed on log-transformed oil values.

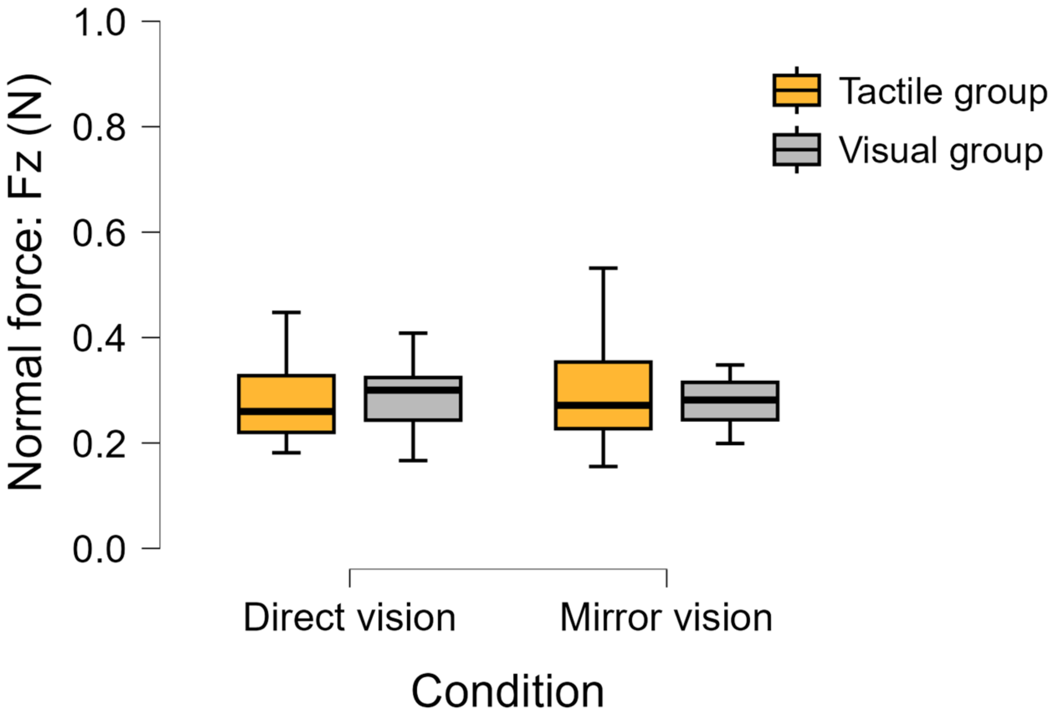

Fig. S2 | **Normal force was stable in both groups during direct and mirror vision conditions.** Boxplots show the normal force (Fz) applied to the surface during the tracing phase, averaged across all trials, for the Tactile (orange) and Visual (grey) groups under Direct vision and Mirror vision conditions. Central lines indicate the median; boxes represent the interquartile range (IQR), and whiskers extend to 1.5× the IQR.

*Table S18. **Normal force (Fz):** Within & between subjects effects.*

| Effect | Sum of squares | df | Mean Square | F | p | $\eta^2 p$ |
|---|---|---|---|---|---|---|
| **Intercept** | 22.8 | 1 | 22.8 | 1042.8 | 0.00 | 22.8 |
| **Group** | 0.00002 | 1 | 0.00002 | 0.001 | 0.98 | 0.00002 |
| **Error** | 0.77 | 35 | 0.02 | | | 0.77 |
| **Condition** | 0.0009 | 1 | 0.0009 | 0.25 | 0.62 | 0.0009 |
| **Group * Condition** | 0.006 | 1 | 0.006 | 1.62 | 0.21 | 0.006 |
| **Error** | 0.13 | 35 | 0.004 | | | 0.13 |

**Table S18** Summary of the 2 (Group: Tactile, Visual) × 2 (Condition: Direct vision, Mirror vision) mixed ANOVA, with repeated measures on the factor Condition on the mean normal force (Fz) applied to the surface during the tracing period. The analysis revealed no significant main or interaction effects, indicating similar normal force between groups and experimental conditions. All analyses were performed on log-transformed values.
